# Critical role of miR-10b in BRaf^V600E^ dependent anchorage-independent growth and invasion of melanoma cells

**DOI:** 10.1101/413476

**Authors:** Ila Datar, Gardiyawasam Kalpana, Ivana de la Serna, Robert Trumbly, Jungmin Choi, Kam C. Yeung

## Abstract

Recent high-throughput-sequencing of cancer genomes has identified oncogenic mutations in the *BRaf* genetic locus as one of the critical events in melanomagenesis. *BRaf* encodes a serine/threonine kinase that regulates the MAPK/ERK kinase (MEK) and extracellular signal-regulated kinase (ERK) protein kinase cascade. In normal cells, the activity of BRaf is tightly regulated and is required for cell growth and survival. *BRaf* gain-of-function mutations in melanoma frequently lead to unrestrained growth, enhanced cell invasion and increased viability of cancer cells. Although it is clear that the invasive phenotypes of *BRaf* mutated melanoma cells are stringently dependent on BRaf-MEK-ERK activation, the downstream effector targets that are required for oncogenic BRaf-mediated melanomagenesis are not well defined. miRNAs have regulatory functions towards the expression of genes that are important in carcinogenesis. We observed that miR-10b expression correlates with the presence of the oncogenic *BRaf* (*BRaf*^*V600E*^) mutation in melanoma cells. While expression of miR-10b enhances anchorage-independent growth of *BRaf* wild-type melanoma cells, miR-10b silencing decreases *BRaf*^*V600E*^ cancer cell invasion *in vitro*. Importantly, the expression of miR-10b is required for BRaf^V600E^-mediated anchorage-independent growth and invasion of melanoma cells *in vitro*. Taken together our results suggest that miR-10b is an important mediator of oncogenic BRaf^V600E^ activity in melanoma.

## Introduction

Melanoma is the most aggressive of all the skin cancers. BRaf is a serine/threonine protein kinase that activates the MEK/ERK-signaling pathway. About 25-70% of malignant melanomas harbor gain-of-function mutations in the oncogene *BRaf* (1). Among several BRaf gain-of-function mutations, BRaf^V600E^ is the most common mutation and accounts for nearly 80% of them (1). Not only does BRaf^V600E^ cause a sustained activation of ERK signaling pathway in melanoma, it is also critical for the malignant process and is one of the few identified driver mutations essential for melanoma proliferation and survival. The transformation of melanocytes to melanoma by BRaf^V600E^ requires activation of the MEK-ERK kinases cascade with multiple downstream components (2-4). The mechanism that integrates the diverse components into a coordinated response to the BRaf^V600E^ mutation remains undefined.

MicroRNAs (miRNAs) are small, non-protein coding RNA molecules and they regulate gene expression through a combination of translational repression and mRNA destabilization (5). Each miRNA targets ∼200 mRNA molecules (6). Because of their pleiotropic potentials, miRNAs are attractive candidates as master regulators of the BRaf^V600E^ oncogenic transformation program. In this study, we identified for the first time, significant positive correlation between BRaf^V600E^ mutation and microRNA-10b (miR-10b) expression. Furthermore, we show that miR-10b is a novel downstream effector of BRaf^V600E^ and that BRaf^V600E^ plays a causal role in the induction of miR-10b in melanoma cell lines. We also show that miR-10b induced by BRaf^V600E^ is able to increase invasive capacity and anchorage-independent growth of melanoma cells.

## Materials and methods

### Cell culture

All cell culture media were from HyClone. Fetal bovine serum (FBS) was from Atlanta Biologicals, and newborn calf serum was from Lonza. 10 cm^2^ and 6 cm^2^ cell culture plates were from Sarstedt. All the melanoma cell lines were cultured in Dulbecco’s modified Eagle’s medium with 10% FBS and 1% penicillin streptomycin and were grown in a humidified tissue culture incubator at 37°C in 5% CO2. Mel 505 (7), PMWK (8), SK-MEL 28 (9), SK-MEL 24 (7), VMM39 (7) and MEL 224 (7) cells were kindly provided by Dr. J. Shields (University of North Carolina, Chapel Hill). YUHEF (10) and YUROB (10) cells were a kind gift from Dr. R. Halaban (Yale University, Connecticut). SK-MEL 197 (11) cells were received from Memorial Sloan Kettering Cancer Center. M249 (12) and M238 (12) cells were a kind gift from Dr. A. Ribas (UCLA).

### Chemicals and reagents

PLX4032 (Vemurafenib) and 4-OHT (4-hydroxy tamoxifen) were purchased from Selleck chemicals and Sigma, respectively. ERK (1:1000), phospho ERK (1:1000), BRaf primary antibodies (1:1000) and horseradish peroxidase secondary antibodies were from Santa Cruz Biotechnology, and tubulin (1:1000) antibody was from Sigma.

### Ectopic expression and depletion of miR-10b

Mammalian expression vectors, MDH1-PGK-GFP 2.0-miR-10b and pBABE-puro-miR-10b sponge, were purchased from Addgene (Plasmids were deposited by Weinberg lab) (13). Melanoma cells were stably transduced with viral particles expressing miR-10b or miR-10 sponge as described previously (14).

### *In vitro* Matrigel invasion assay

The polycarbonate membrane (8µ pore size) of FluoroBlok cell culture inserts (BD Biosciences) was coated with 60 uL of Matrigel (1:26 in serum-free medium) (BD Biosciences) and incubated at 37°C for 2-3 hours. 1×10^4^ of SK-Mel 28 cells were plated on these inserts. 700mL of chemo-attractive medium (Dulbecco’s Modified Eagle’s medium, 1% P/S and 10% FBS) was added to the lower chambers (24-well BD Falcon TC companion plate). After 24 hours of incubation, the insert bottoms were dipped in 1X PBS and stained in Calcein AM reconstituted in DMSO to 1mg/mL-1ml in 700 ml 1X PBS (BD Biosciences). Digital images were captured on EVOS inverted microscope along with manual cell count and fluorescence reading (485-538nM).

### RNA extraction and TaqMan miRNA quantitative real-time PCR assay

RNA extraction was carried out using Qiazol reagent (Qiagen). TaqMan miRNA qRT-PCR was performed according to manufacturer’s instructions (TaqMan^®^). RNU6 was used as the internal control. Briefly, in the reverse transcription (RT) step, cDNA was reverse transcribed from total RNA samples using specific looped miRNA primers from the TaqMan miRNA assays and reagents from the TaqMan. In the PCR step, PCR products were amplified from cDNA samples using the TaqMan miRNA assay together with the TaqMan^®^ universal PCR master mix.

### Soft agar assay for anchorage-independence

1mL layer of 1:1 mixture of 1.2% agar and 2X medium (to give a final 0.6% agar concentration) was evenly spread on 35mm plates. It was allowed to set for 30 minutes at RT. Cells were counted and plated in a mixture of 1.2% agar and 2X medium (1:4) to give a final agar concentration of 0.3%. 3×10^3^ of SK-Mel 28 cells or 1×10^4^ of Mel 505 cells or 1×10^4^ of SK-Mel 197 cells were plated. Number of colonies formed were stained with MTT and counted under bright field microscope at 4X objective. Cells were plated in triplicates and four fields were counted per 35 mm plate.

### TCGA data analysis

The publicly available TCGA data were directly downloaded from the TCGA Data Portal at https://tcga-data.nci.nih.gov/. TCGA melanoma cases (TCGA-SKCM) annotated for BRAF WT or V600E mutation were extracted from the GDC portal (https://portal.gdc.cancer.gov). Among those, 407 samples were profiled for both mRNA expression and miRNA expression. Fragments per Kilobase per Million mapped reads (FPKM) from the RNA-sequencing data and reads per million miRNA mapped from the miRNA data were used as normalized read counts. Pearson correlation coefficients and P-values for miR-10b expression with TWIST1 and MYC expression were calculated, respectively.

### Western blot analysis

Cell extracts were prepared and western blotting was carried out. Briefly, cells were lysed with 20 mM Tris, pH 7.4, 150mM NaCl, 2mM EDTA, and 1% Triton X-100. Samples (10–50 μg protein) were separated by sodium dodecyl sulfate-polyacrylamide electrophoresis and then electrophoretically transferred from the gel to polyvinylidene fluoride membranes (Millipore). All the primary antibodies were diluted in phosphate buffered saline, pH 7.4, 0.2% Tween-20, 5% bovine serum albumin and 0.002% sodium azide. Following three washes, blots were incubated with the appropriate horseradish peroxidase-conjugated secondary antibody for 1 h at room temperature. Proteins were visualized with enhanced chemiluminescence with the BioRad ChemiDoc EQ system.

### Statistical analysis

All experiments were performed in triplicates. All the statistical analyses were performed using the two-tailed student’s t-test (GraphPad Prism software).

## Results and discussion

We previously reported that the expression of several miRNAs was altered upon introduction of BRaf^V600E^ in primary melanocytes (15). Among the miRNAs whose expression levels were affected by BRaf^V600E^, we chose miR-10b for further study because its expression positively correlates with V600E mutation in BRaf in established melanoma cell lines (Fig 1), clinical melanoma samples (S2 Fig), and because it is involved in several other malignancies (16).

**Fig 1.**
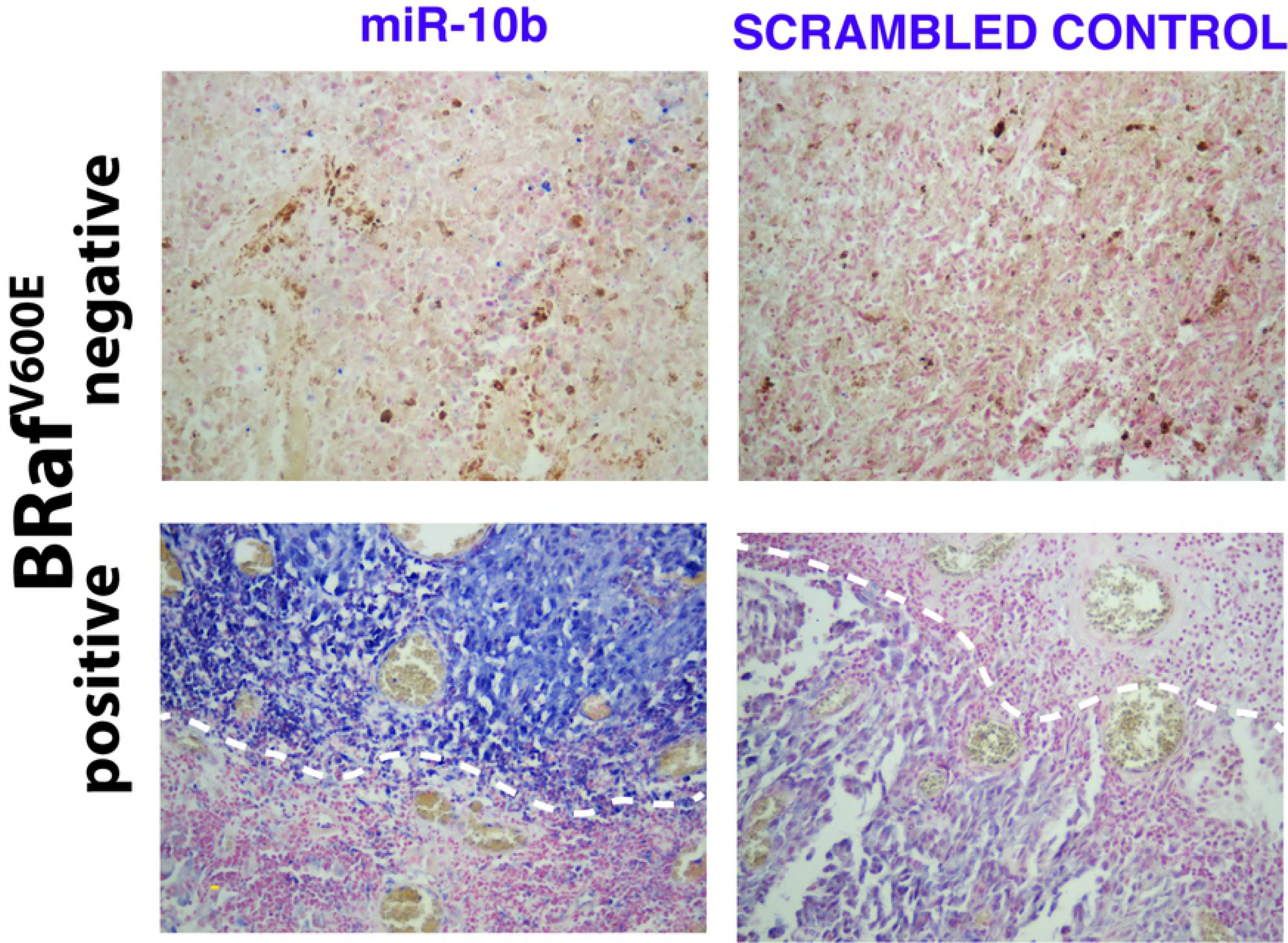
miR-10b expression positively correlates with BRaf^V600E^ mutation status in multiple cell lines. TaqMan reverse transcriptase PCR shows a strong positive correlation of miR-10b expression with BRaf^V600E^ or N-Ras activating mutations in a panel of melanoma cell lines. The experiment was repeated 3X with similar results.

To investigate if BRaf^V600E^ plays a causal role in regulating miR-10b expression in melanoma cell lines we undertook a loss-of-function approach. We stably knocked down BRaf^V600E^ expression by siRNA in two different BRaf^V600E^ melanoma cell lines. As expected, BRaf protein expression significantly decreased in BRaf^V600E^ knockdown cells with a concomitant decrease in phosphorylation of ERK1/2. In both studied cell lines, we found that miR-10b expression was significantly down-regulated upon BRaf^V600E^ knock down (Fig 2a). A decrease in miR-10b expression was also observed in M249, which are BRaf^V600E^ positive melanoma cells when BRaf^V600E^ activity was down-regulated by vemurafenib (PLX4032), a pharmacological inhibitor of BRaf^V600E^ (Fig 2b).

**Fig 2.**
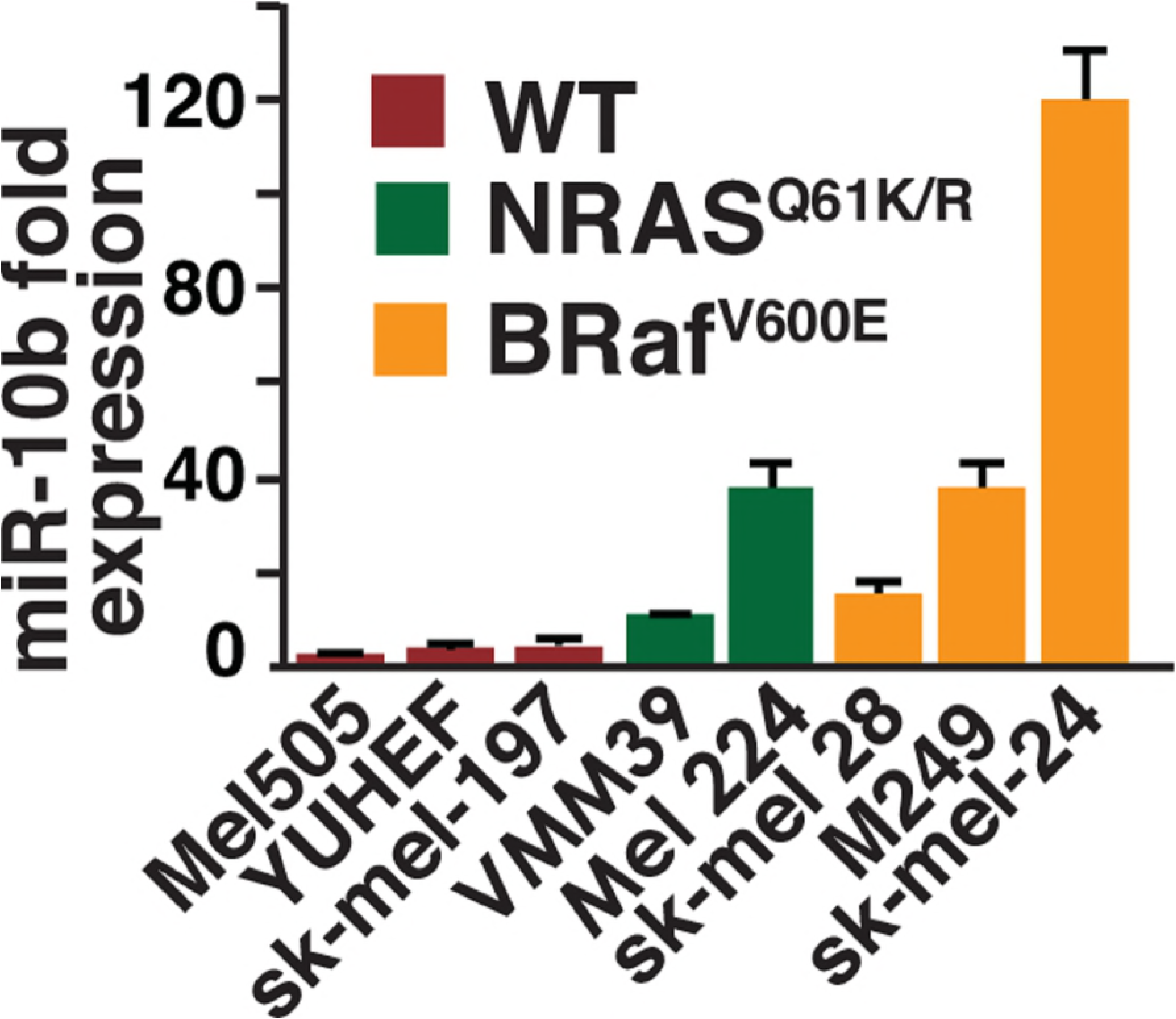
BRaf^V600E^ plays a causal role in the induction of miR-10b. (a) Lower panel shows downregulation of miR-10b expression upon stable knock-down of BRaf^V600E^ in melanoma cell lines SK-MEL 24 and SK-MEL 28 with BRaf^V600E^-specific siRNA as determined by TaqMan qRT-PCR. Upper panel: Western blots of lysates from cells expressing siBRaf^V600E^ or siGFP siRNA. (b) TaqMan qRT-PCR shows downregulation of miR-10b upon inhibition of the BRaf^V600E^ in melanoma cell line M249 by PLX4032. Cells were treated at 1µM concentration of PLX4032 (Vemurafenib) or DMSO for a period of 12 hours. (c) Increase in the expression of miR-10b upon ectopic expression of BRaf^V600E^. Cells were stably transduced with BRaf^V600E^ - or empty vector-expressing retroviruses. (d) Potent induction of miR-10b in Mel 505 cell line that carries wild type (WT) BRaf, upon induction of BRaf^V600E^ following 24 hours 4-OHT (4-hydroxy tamoxifen) treatment at 1µM concentration. Cells were stably transduced with BRaf^V600E^:ERTM-expressing retrovirus.

We reasoned that if the effect of knocking down of BRaf^V600E^ expression on miR-10b is physiological we should observe an opposite effect with over-expression of the oncogenic BRaf^V600E^ in melanoma cell lines carrying wild-type BRaf. Indeed, we found that the expression of miR-10b was significantly upregulated in cells ectopically expressing BRaf^V600E^ as compared to empty vector control (Fig 2c). The drawback of over-expression of an oncogene is that the cells might get adapted to its sustained expression and this can lead to non-specific effects that may not be attributable to the expression of the oncogene alone. As a complementary approach, we used an estrogen receptor fusion system where BRaf^V600E^ was rendered functionally hormone-dependent by fusion with synthetic steroid 4-hydroxytamoxifen (4-OHT)-binding domain of mutated estrogen receptor (ER^T2^) (17). As expected, upon treating the cells with 4-OHT for 24 hours, we observed potent induction of ERK dual phosphorylation at Thr^202^ and Tyr^204^ indicating that the kinase activity of BRaf^V600E^ was temporally turned on by 4-OHT treatment (Fig 2d). Interestingly, transient activation of BRaf^V600E^ is better at activating miR-10b expression than sustained expression of the constitutively active kinase (Compare Figs 2c and 2d). Taken together, our results suggest that the expression of miR-10b in melanoma is BRaf^V600E^ dependent.

At present, it is not clear how the expression of miR-10b is regulated by BRaf^V600E^ in melanoma cells. In humans, the *miR-10b* gene resides upstream from the developmental gene *Hoxd4*. It has been shown that the basic helix-loop-helix transcription factor Twist positively regulates miR-10b expression in breast cancer cells by directly binding to the proximal E-box present in the *miR-10b* gene promoter. The expression levels of Twist are often deregulated in a vast majority of melanoma tumors and cell lines (18, 19). Furthermore, the expression of Twist was shown to be regulated at the transcription level by BRaf^V600E^ mutation in human melanoma cells (20), raising the possibility that BRaf^V600E^ may increase miR-10b expression by activating Twist. Indeed, we observed significant correlation of miR-10b and Twist1 expression only in BRaf^V600E^ mutated melanoma clinical samples (Fig 3a and Table 1). The observed correlation is specific as no significant correlation was observed between another E-box transcription factor c-myc and miR-10b (Fig 3b and Table 2). Further studies validating the promoter occupancy of miR-10b by Twist in the context of melanoma are thus warranted.

**Table 1.**
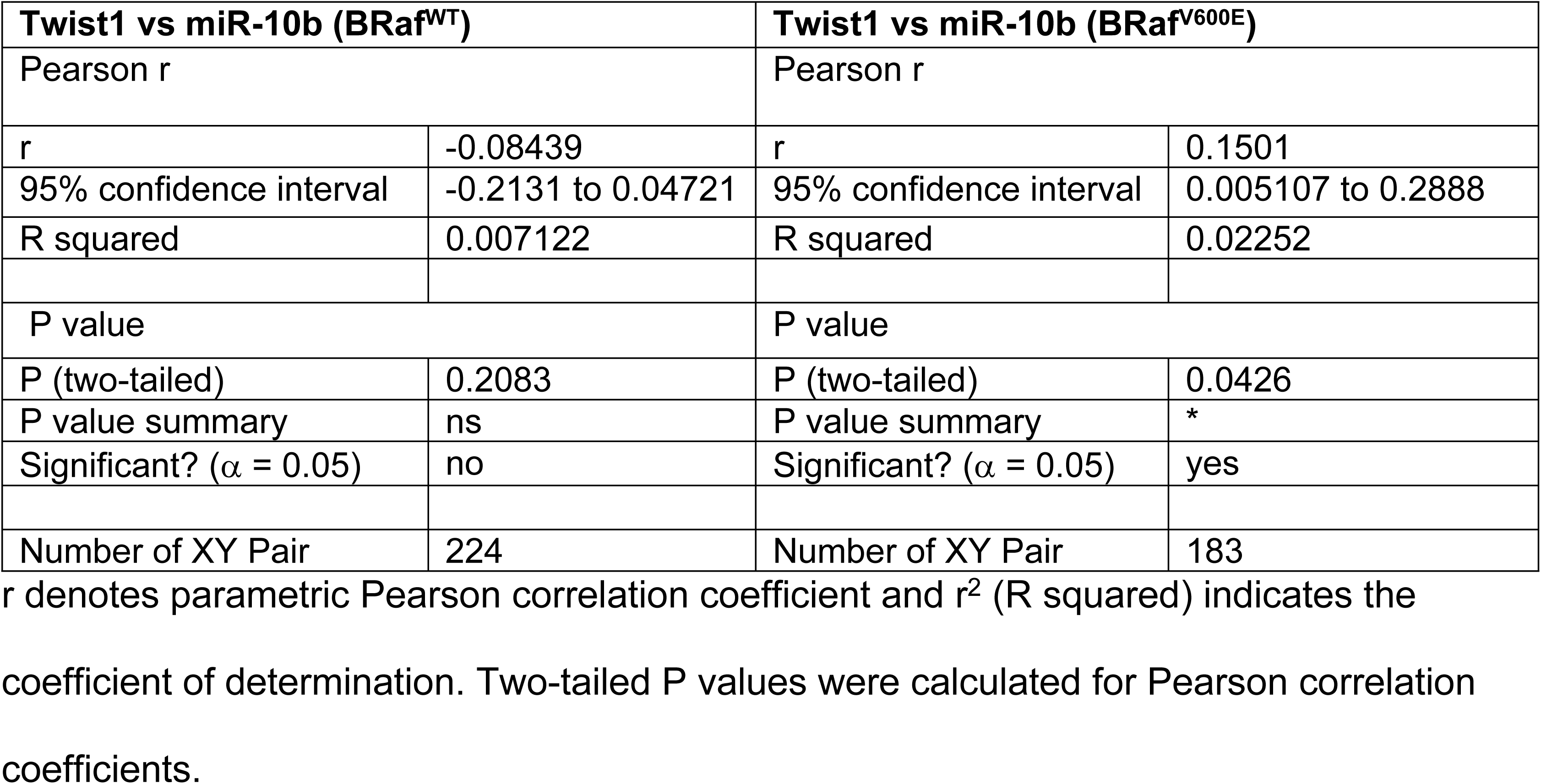
Pearson correlation statistics between Twist1 and miR-10b expression in BRaf^WT^ and BRaf^V600E^.

**Table 2.** Pearson correlation statistics between Myc and miR-10b expression in BRaf^WT^ and BRaf^V600E^.

| c-Myc vs miR-10b (BRAF<sup>WT</sup>) |  | c-Myc vs miR-10b (BRAF<sup>V600E</sup>) |  |
| --- | --- | --- | --- |
| Pearson r |  | Pearson r |  |
| r | 0.08854 | r | -0.03904 |
| 95% confidence interval | -0.04304 to 0.2171 | 95% confidence interval | -0.1831 to 0.1066 |
| R squared | 0.007839 | R squared | 0.001524 |
| P value |  | P value |  |
| P (two-tailed) | 0.1867 | P (two-tailed) | 0.5998 |
| P value summary | ns | P value summary | ns |
| Significant? ($\alpha = 0.05$) | no | Significant? ($\alpha = 0.05$) | no |
| Number of XY Pair | 224 | Number of XY Pair | 183 |

r denotes parametric Pearson correlation coefficient and $r^2$ (R squared) indicates the coefficient of determination. Two-tailed P values were calculated for Pearson correlation coefficients.

**Fig 3.**
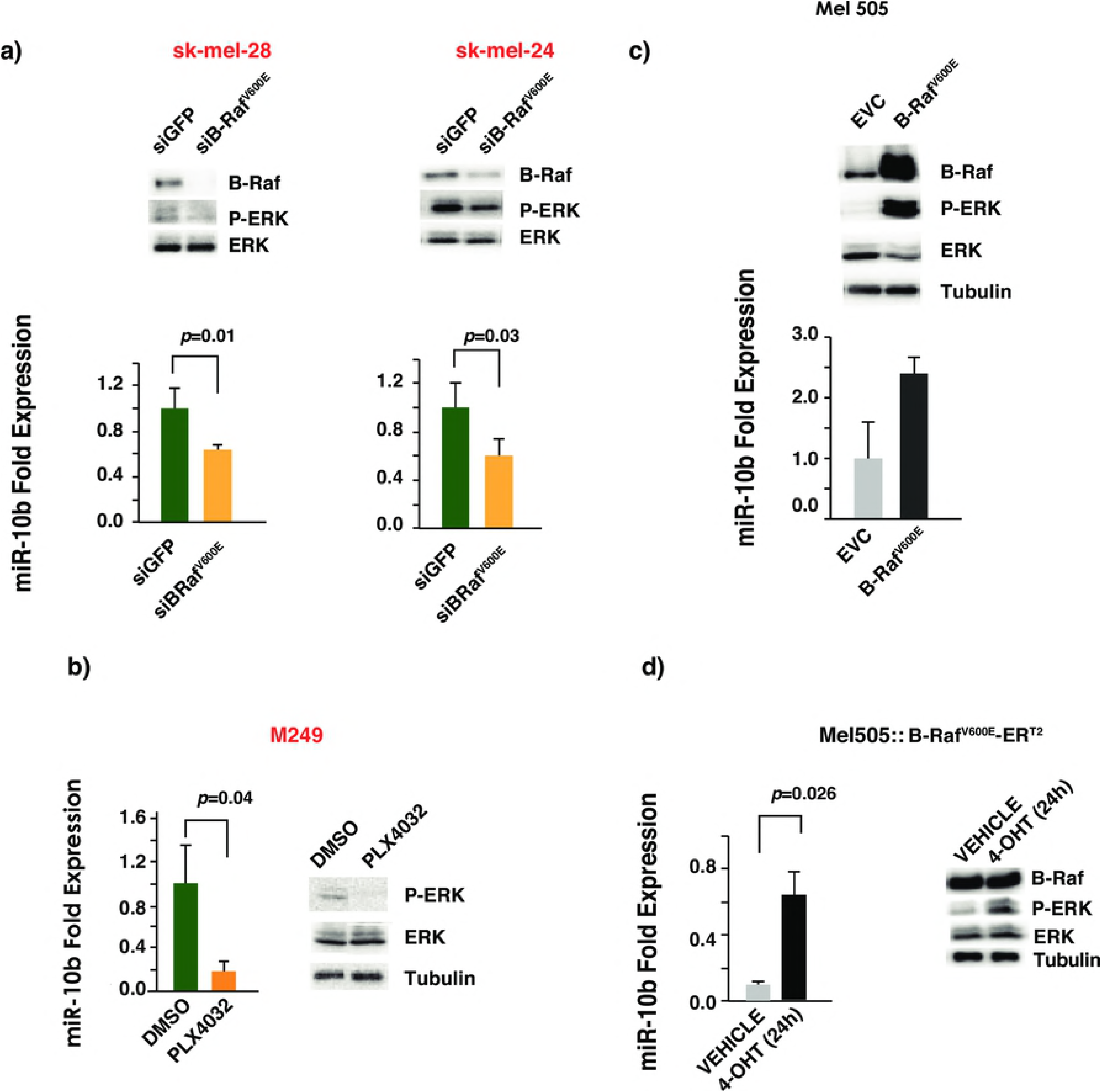
miR-10b displays significant correlation with Twist1 mRNA expression in BRaf^V600E^ mutant melanoma. Scatter plots showing correlation of miR-10b expression with (a-b) Twist1 mRNA expression and (c-d) Myc mRNA expression in BRAF^WT^ and BRaf^V600E^ samples from 407 TCGA-SKCM samples. r denotes Pearson correlation coefficient. p refers to a Pearson correlation P-value.

Studies with cancer cell transplantation and autochthonous cancer mouse models have demonstrated the causal role of miR-10b in breast cancer initiation, growth, progression and metastasis (21). However, the causality of loss- or gain-of-function of miR-10b in other cancers including melanoma is not known. Anchorage-independent growth and increased invasive capacity are two well-characterized hallmarks of cancer. miR-10b may also be involved in conferring these important cancerous phenotypes on melanoma cells. Indeed, in low-miR-10b-expressing wild-type *BRaf* melanoma cells Mel 505, ectopic expression of miR-10b was sufficient to increase anchorage-independent growth as measured in soft agar (Fig 4a). This effect was not cell type specific as a similar effect was observed with other wild-type *BRaf* melanoma cells SK-MEL197. Similarly, depletion of miR-10b in high-miR-10b-expressing *BRaf*^*V600E*^ melanoma cells, SK-MEL 28, decreased their invasiveness (Fig 4b). BRaf^V600E^ is an oncoprotein and its presence is essential for anchorage-independent growth and invasive capacity of *BRaf*^*V600E*^ melanoma cells. BRaf^V600E^ induces miR-10b expression in melanoma cell lines. Therefore, *BRaf*^*V600E*^ melanoma cells may acquire cancer phenotypes partly by increasing the expression levels of miR-10b. Consistent with this line of thinking, ectopic miR-10b expression significantly rescued the decrease in anchorage-independent growth and invasiveness due to the loss of BRaf^V600E^ in SK-MEL 28 cells (Fig 5).

**Fig 4.**
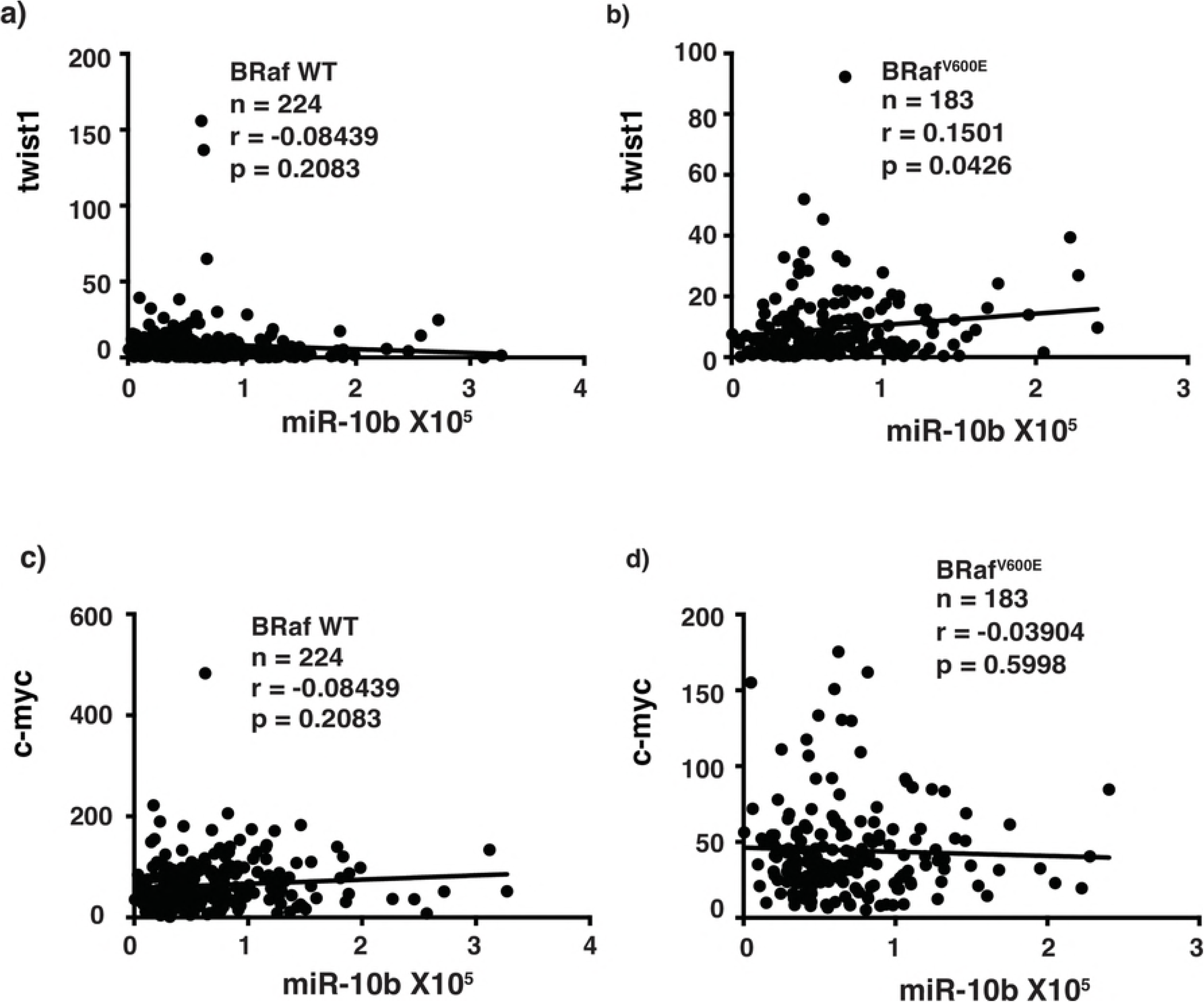
miR-10b confers anchorage-independence and increases invasive capacity of melanoma cell lines. (a) Lower panel shows increase in the number of colonies formed in soft agar upon ectopic expression of miR-10b in Mel 505 and SK-MEL 197 melanoma cell lines expressing WT BRaf. Upper panel shows miR-10b fold expression upon ectopic expression of miR-10b. Student’s t-test was performed for statistical analysis, which shows a significant value of 0.002 for Mel 505 and 0.005 for SK-MEL 197 cell lines. (b) Lower left panel shows decrease in number of invading cells upon miR-10b depletion by specific miR-10b sponge. Lower right panel shows a representative field of Matrigel membrane with invaded cells stained fluorescent green with Calcein AM. Student’s t-test was performed and a significant difference was found with a p-value of 0.0001. Upper panel shows expression levels of miR-10b in miR-10b sponge expressed SK-MEL 28 cells as determined by TaqMan qRT-PCR.

**Fig 5.**
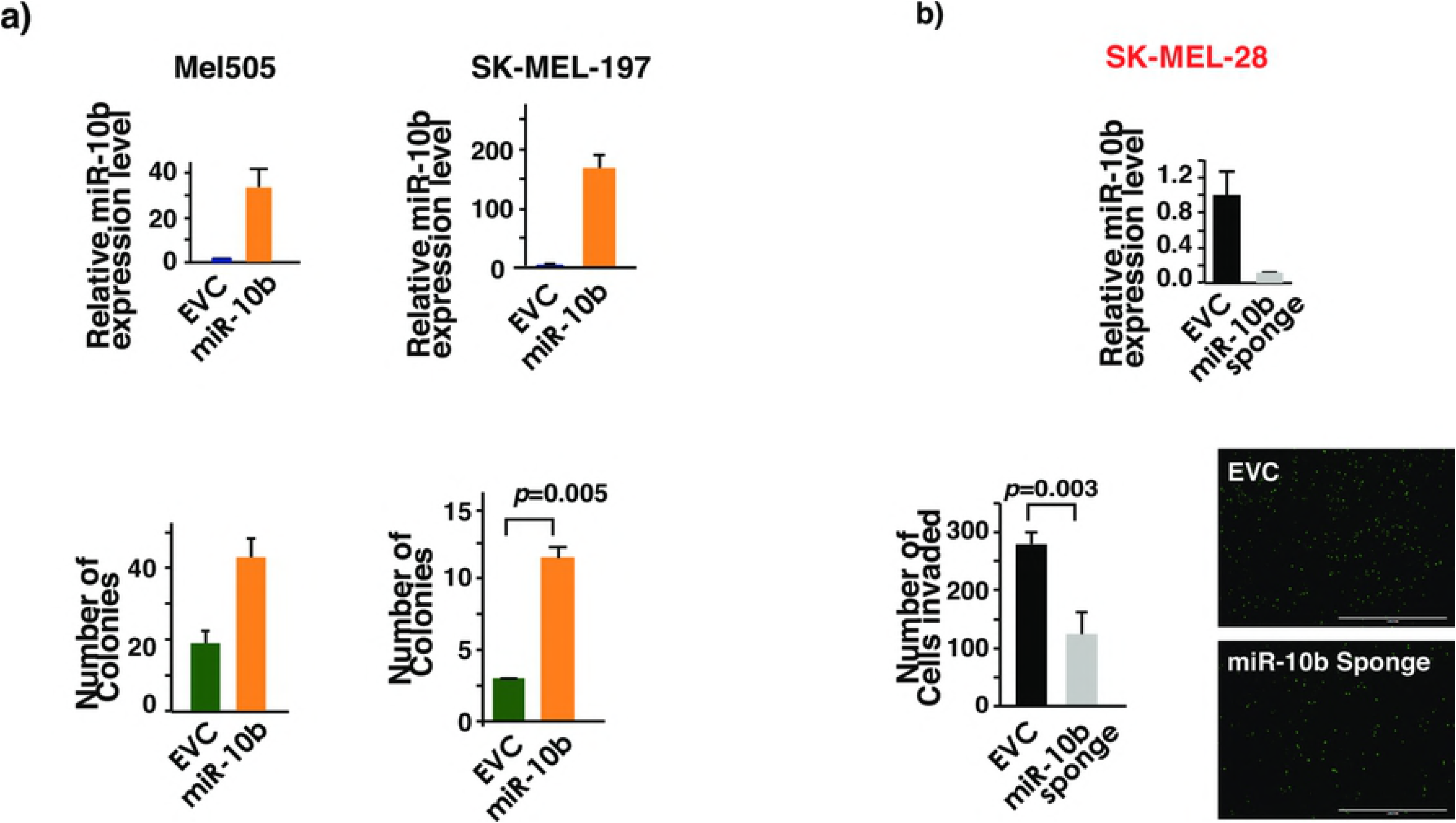
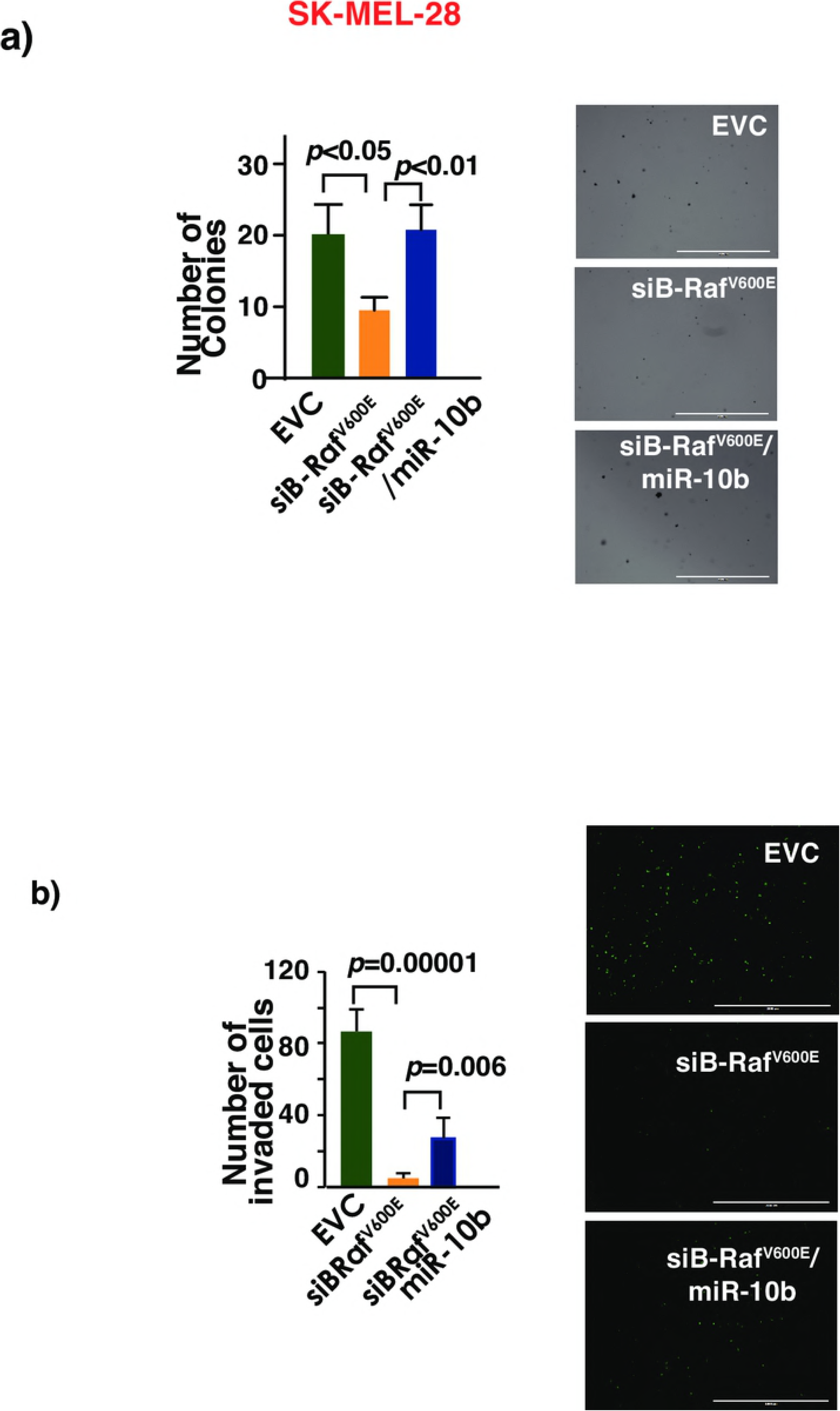
BRaf^V600E^ melanoma cells acquire cancer phenotypes partly by increasing the expression levels of miR-10b. (a) Upper panel: BRaf^V600E^ knockdown decreases number of colonies in soft agar in SK-MEL 28 cell line. Restoration of miR-10b expression reversed this effect. This experiment was performed at least twice with triplicates each time. ANOVA was performed followed by Bonferroni’s multiple comparison test where EVC vs siBRaf^V600E^ and BRaf^V600E^ vs BRaf^V600E^ /miR-10b were found to be significant with a p-value <0.05. EVC-empty vector control. (b) Knockdown of BRaf^V600E^ decreases melanoma cell invasion while restoration of miR-10b reverses this effect. Right panel shows a representative field of Matrigel membrane with invading cells staining fluorescent green with Calcein AM. One-way ANOVA was performed with p-value<0.05.

The mechanisms of how augmented miR-10b expression in BRaf^V600E^ melanoma cells enhances their anchorage-independent growth and invasiveness is currently not known. miRNAs function mainly by silencing expression of target genes by binding to specific sites of the targeted gene mRNAs. Studies of genetic deletion of miR-10b in MMTV-PyMT mice identified *Tbx5, Pten*, and *HoxD10* as key miR-10b targets (21). While *Tbx5* and *Pten* are well-characterized tumor suppressor genes, *HoxD10* is a proven metastasis suppressor gene. To grow continuously after being detached from extracellular matrix, cancer cells develop resistance to a specific form of caspase-mediated cell death known as anoikis by upregulating the survival pathways. PI3K pathway is a major survival pathway cancer cells employ to overcome anoikis to achieve anchorage-independent growth (22). Of the three target genes of miR-10b, *Pten* has been shown to play a pivotal role in dampening the PI3K pathway. It is therefore possible that miR10b enhances anchorage-independent growth in melanoma by decreasing the expression of Pten. Both Tbx2 and HoxD10 are transcription factors that can potentially regulate a multitude of genes. It has been shown that HoxD10 represses the expression of metastasis gene *RhoC* in breast cancer cells and that miR-10b increases cancer cells invasion by repressing the repressor of *RhoC* expression. *RhoC* is a well-studied metastasis gene in melanoma and it is possible that BRaf^V600E^ utilizes the same miR-10b-HoxD10-RhoC axis to drive cancer cells invasion. Likewise, Tbx2 can also play an equally important role in mediating the different effects on melanomagenesis resulting from BRaf mutation.

In 2010, an orally available inhibitor of BRaf^V600E^ increased the median progression-free survival of patients with malignant melanoma harboring this mutation, to more than 7 months (23). However, patients develop resistance to this drug due to several mechanisms including receptor tyrosine kinase (RTK) or N-Ras upregulation among many others (24). Recently, combinatorial therapy utilizing BRaf and MEK inhibition delayed the emergence of resistance while progression-free survival remained more or less similar to BRAF inhibition alone (25). However, relapse due to drug-resistance is commonplace and warrants a treatment strategy based on new downstream molecular targets of the mutant BRaf protein. The expression of miR-10b is often deregulated in cancers including melanoma and its expression was positively associated with worse patient survival (26). Here we identify miR-10b as a novel molecular target downstream of mutant BRaf protein raising the possibility that targeting miR-10b and/or its effectors may represent a rational alternative treatment for relapsed patient with BRaf^V600E^ melanoma.

## Acknowledgements

UT Foundation, a gift from Clement Lam to KCY, supported this work.

## Supporting information

**S1 Method. miRNA *in situ* hybridization**

**S2 Fig. Expression levels of miR-10b in clinical melanoma samples as determined by ISH with labeled specific miRNA probe.** (n= 3). The BRaf^V600E^ mutation was determined by IHC staining with BRaf^V600E^ specific Ab. Lesions were demarcated from normal tissues with white dotted line.

## References

1. Garnett MJ, Marais R. Guilty as charged: B-RAF is a human oncogene. Cancer Cell. 2004;6(4):313–9.

2. Acosta JC, O’Loghlen A, Banito A, Guijarro MV, Augert A, Raguz S, et al. Chemokine signaling via the CXCR2 receptor reinforces senescence. Cell. 2008;133(6):1006–18.

3. Kuilman T, Michaloglou C, Vredeveld LC, Douma S, van Doorn R, Desmet CJ, et al. Oncogene-induced senescence relayed by an interleukin-dependent inflammatory network. Cell. 2008;133(6):1019–31.

4. Wajapeyee N, Serra RW, Zhu X, Mahalingam M, Green MR. Oncogenic BRAF induces senescence and apoptosis through pathways mediated by the secreted protein IGFBP7. Cell. 2008;132(3):363–74.

5. Eichhorn SW, Guo H, McGeary SE, Rodriguez-Mias RA, Shin C, Baek D, et al. mRNA destabilization is the dominant effect of mammalian microRNAs by the time substantial repression ensues. Mol Cell. 2014;56(1):104–15.

6. He L, Hannon GJ. MicroRNAs: small RNAs with a big role in gene regulation. Nat Rev Genet. 2004;5(7):522–31.

7. Park S, Yeung ML, Beach S, Shields JM, Yeung KC. RKIP downregulates B-Raf kinase activity in melanoma cancer cells. Oncogene. 2005;24:3535.

8. Vink J, Thomas L, Etoh T, Bruijn JA, Mihm MC, Jr., Gattoni-Celli S, et al. Role of beta-1 integrins in organ specific adhesion of melanoma cells *in vitro*. Lab Invest. 1993;68(2):192–203.

9. Zhang XD, Franco A, Myers K, Gray C, Nguyen T, Hersey P. Relation of TNF-related apoptosis-inducing ligand (TRAIL) receptor and FLICE-inhibitory protein expression to TRAIL-induced apoptosis of melanoma. Cancer Res. 1999;59(11):2747– 53.

10. Halaban R, Zhang W, Bacchiocchi A, Cheng E, Parisi F, Ariyan S, et al. PLX4032, a selective BRAFV600E kinase inhibitor, activates the ERK pathway and enhances cell migration and proliferation of BRAF melanoma cells. Pigment cell & melanoma research. 2010;23(2):190–200.

11. Kelly KJ, Brader P, Woo Y, Li S, Chen N, Yu YA, et al. Real-time intraoperative detection of melanoma lymph node metastases using recombinant vaccinia virus GLV-1h68 in an immunocompetent animal model. International journal of cancer. 2009;124(4):911–8.

12. Søndergaard JN, Nazarian R, Wang Q, Guo D, Hsueh T, Mok S, et al. Differential sensitivity of melanoma cell lines with BRAF V600E mutation to the specific Raf inhibitor PLX4032. Journal of Translational Medicine. 2010;8(1):39.

13. Ma L, Teruya-Feldstein J, Weinberg RA. Tumour invasion and metastasis initiated by microRNA-10b in breast cancer. Nature. 2007;449(7163):682–8.

14. Ma J, Shi J, Zhao D, Cheng L, Wang W, Li F, et al. Raf kinase inhibitor protein inhibits cholangiocarcinoma cell metastasis by downregulating matrix metalloproteinase 9 and upregulating tissue inhibitor of metalloproteinase 4 expression. Oncology letters. 2015;9(1):15–24.

15. Ren G, Feng J, Datar I, Yeung AH, Saladi SV, Feng Y, et al. A Micro-RNA Connection in BRafV600E-Mediated Premature Senescence of Human Melanocytes. Int J Cell Biol. 2012;2012:913242.

16. Lund AH. miR-10 in development and cancer. Cell Death Differ. 2010;17(2):209– 14.

17. Littlewood TD, Hancock DC, Danielian PS, Parker MG, Evan GI. A modified oestrogen receptor ligand-binding domain as an improved switch for the regulation of heterologous proteins. Nucleic Acids Res. 1995;23:1686–90.

18. Ansieau S, Bastid J, Doreau A, Morel AP, Bouchet BP, Thomas C, et al. Induction of EMT by twist proteins as a collateral effect of tumor-promoting inactivation of premature senescence. Cancer Cell. 2008;14(1):79–89.

19. Hoek K, Rimm DL, Williams KR, Zhao H, Ariyan S, Lin A, et al. Expression profiling reveals novel pathways in the transformation of melanocytes to melanomas. Cancer Res. 2004;64(15):5270–82.

20. Weiss MB, Abel EV, Mayberry MM, Basile KJ, Berger AC, Aplin AE. TWIST1 is an ERK1/2 effector that promotes invasion and regulates MMP-1 expression in human melanoma cells. Cancer Res. 2012;72(24):6382–92.

21. Kim J, Siverly AN, Chen D, Wang M, Yuan Y, Wang Y, et al. Ablation of miR-10b Suppresses Oncogene-Induced Mammary Tumorigenesis and Metastasis and Reactivates Tumor-Suppressive Pathways. Cancer Res. 2016;76(21):6424–35.

22. Paoli P, Giannoni E, Chiarugi P. Anoikis molecular pathways and its role in cancer progression. Biochim Biophys Acta. 2013;1833(12):3481–98.

23. Flaherty KT, Puzanov I, Kim KB, Ribas A, McArthur GA, Sosman JA, et al. Inhibition of mutated, activated BRAF in metastatic melanoma. N Engl J Med. 2010;363(9):809–19.

24. Nazarian R, Shi H, Wang Q, Kong X, Koya RC, Lee H, et al. Melanomas acquire resistance to BRAFV600E inhibition by RTK or N-RAS upregulation. Nature. 2010;468(7326):973–7.

25. Long GV, Stroyakovskiy D, Gogas H, Levchenko E, de Braud F, Larkin J, et al. Combined BRAF and MEK inhibition versus BRAF inhibition alone in melanoma. N Engl J Med. 2014;371(20):1877–88.

26. Saldanha G, Elshaw S, Sachs P, Alharbi H, Shah P, Jothi A, et al. microRNA-10b is a prognostic biomarker for melanoma. Mod Pathol. 2016;29(2):112–21.

